# The miR-183/96/182 Cluster Regulates Trigeminal Ganglion Sensory Neurons’ Response to *Pseudomonas aeruginosa* Infection

**DOI:** 10.64898/2026.03.30.715374

**Authors:** Giovanni LoGrasso, Naman Gupta, Sai Giridhar Reddy Bugulu, Linda D. Hazlett, Anthony J. St. Leger, Shunbin Xu

**Affiliations:** Department of Ophthalmology, Visual and Anatomical Sciences, Wayne State University School of Medicine/Kresge Eye Institute, Detroit, Michigan; Department of Ophthalmology, University of Pittsburgh, Pittsburgh, Pennsylvania, United States; Immunology, University of Pittsburgh, Pittsburgh, Pennsylvania, United States

## Abstract

**Purpose:** To uncover the molecular mechanisms of corneal sensory nerves (CSN)’s involvement in the initiation of *Pseudomonas aeruginosa* (PA) keratitis and the roles of the miR-183/96/182 cluster (miR-183C) in this process.

**Methods:** miR-183C conventional knockout (KO) or sensory neuron-specific (SNS) conditional (C)KO mice and their age- and sex-matched wild type (WT) controls were used. TG SN were isolated. Neurite growth and branching were analyzed by neurite tracing. Custom-made microfluidic chambers (MFC) were used to separate the neuronal cell bodies in the soma chamber and their neurites/nerve endings in the axon chamber. TG SN’s response to lipopolysaccharide (LPS) or PA infection of the neurites/nerve endings was studied by ELISA assays of CX3CL1 and substance P (sP) in the axon chamber. Target luciferase reporter assays were performed to validate key downstream target genes of miR-183C.

**Results:** The total neurite length and number of branches per TG SN were decreased in the CKO vs WT mice, and in the male vs female WT mice. PA infection, but not LPS alone, induced the production and secretion of CX3CL1 and sP in WT mice; while TG SN of miR-183C KO mice responded to both LPS and PA and were significantly enhanced when compared to WT mice. Antagonists to TLR4 and/or FPR1 inhibited PA-induced responses. Target luciferase reporter assays confirmed that genes encoding NRP1, TAC1-the precursor gene of sP, CX3CL1 and ADAM10, a metalloproteinase involved in the production of soluble CX3CL1, were direct targets of miR-183C.

**Conclusions:** PA directly activates TG SN and induces chemokine and neuropeptide production/secretion through TLR4 and FPR1 receptors, which may contribute to the initiation of PA keratitis. miR-183C regulates TG SN neurite growth, chemokine and neuropeptide production/secretion and the response to PA infection by targeting a collection of key genes involved in axon guidance/projection-, chemokine and neuropeptide biogenesis- and receptors mediating PA-induced activation.

## Introduction

*Pseudomonas aeruginosa* (PA) is a Gram-negative bacterium and an opportunistic pathogen. It is a common causative organism associated with ocular injury of agricultural accidents-related bacterial keratitis in developing countries, and contact lens-related disease in developed countries^1, 2^. PA-induced keratitis is one of the most rapidly developing and destructive diseases of the cornea^1^. Currently, PA keratitis is mainly treated by topical administration of antibiotics; in severe cases, subconjunctival injection may be employed. Although antibiotic treatment reduces the bacterial burden, tissue damage often occurs as a result of a poorly controlled host immune response^3, 4^. Additionally, frequent emergence of antibiotic resistant strains poses serious challenges for the effective management of the disease^5–7^. It is urgent and timely to develop alternative treatment approaches.

Currently, most of the studies on the molecular mechanisms of the initiation of PA keratitis have focused on corneal epithelial and immune cells^8, 9^. Numerous elegant studies have shown that corneal epithelial cells can detect PA bacteria through pattern recognition receptors, e.g., TLR4 and TLR5, secrete antimicrobial peptides and initiate an inflammatory response by releasing pro-inflammatory cytokines/chemokines, which further recruit innate and adaptive immune cells from the circulation^1, 8, 9^. These studies provided important insights into the roles of corneal epithelial and immune cells in the pathogenesis of PA keratitis; however, the role of another important component of the corneal niche, corneal sensory nerves (CSN), have not been adequately studied.

The cornea is the most densely innerved tissue in the body^10–12^. Myelinated sensory nerves from the trigeminal ganglion (TG) become unmyelinated while entering the corneal stroma and extend to the center of the cornea. CSN in the stroma enter the epithelial layer by piercing through the epithelial basement membrane. Once through the basement membrane, they change their orientation and extend horizontal to the basement membrane toward the center, forming the whorl-like patterned intra-epithelial corneal basal nerve plexus (ICBNs)^11, 13–16^. The ICBN bundles further branches and extends their terminals apically between epithelial cells as far as the tight junctions formed by the surface squamous epithelial cells^13, 15, 16^. It is estimated that the cornea contains approximately 7000 free epithelial nerve endings/ mm^2^, >80,000 per cornea in human^11, 14, 17^, and ∼40,000-80,000 per cornea in mouse^18–20^. Therefore, we hypothesized that, in addition to corneal epithelial and immune cells, a response by CSN to bacterial infection plays a major role in initiation of PA keratitis. Although increasing evidence suggests that neuroimmune interactions modulate corneal immune response to PA infection^21–31^, whether and how CSN are involved in the initiation of PA keratitis is still unknown.

microRNAs (miRNAs) are small, non-coding RNAs and are important post-transcriptional regulators of gene expression^32–35^. They are proven to play an important role in human diseases^36–43^ and are viable therapeutic targets^44–47^. However, their precise roles in bacterial keratitis remain largely unexplored. Previously, in the exploration of the roles of miRNAs in bacterial keratitis, we identified a conserved, paralogous miRNA cluster, the miR-183/96/182 cluster (miR-183C), which is highly specifically expressed in all primary sensory neurons of all major sensory organs, including the TG, where the sensory neurons innervating the cornea reside^28, 48, 49^. In addition, miR-183C is also expressed in innate immune cells, including macrophages (Mφ) and polymorphonuclear leukocytes (PMN) or neutrophils^28^. Inactivation of miR-183C in a conventional knockout (KO) mouse model results in a decreased CSN density, reduced levels of pro-inflammatory neuropeptides and chemokines/cytokines and decreased severity of PA keratitis^28^. In innate immune cells, miR-183C regulates both their cytokine production and phagocytosis and bacterial killing capacity through targeting key genes involved in these processes^27, 28, 50, 51^. In regard to its functions in sensory innervation, we showed that sensory neuron-specific conditional knockout (SNS-CKO) of miR-183C also results in decreased CSN density^27^; and acute knockdown of miR-183C in the cornea causes a reversible regression of CSN^52^. Data from RNA sequencing in the TG suggests that miR-183C imposes intrinsic regulation of CSN by directly targeting neuronal projection and axon guidance-related genes^27^. Intriguingly, SNS-CKO of miR-183C also results in an increased number of corneal resident myeloid cells in steady-state corneas, suggesting an extrinsic regulation of miR-183C of corneal innate immunity through neuroimmune interaction^27^.

To uncover the mechanisms of how CSN respond to PA infection in the initiation of PA keratitis and the roles of miR-183C in these processes, in this study, we directly characterized the effect of inactivation of miR-183C on neurite growth of TG SN and their responses to PA infection. In addition, we identified and validated several predicted target genes known to play key roles in these processes. Here we provide evidence that PA induces the production and secretion of neuron-produced chemokine CX3CL1 and neuropeptide substance P (sP), through receptor molecules, TLR4 and FPR1, on TG SN neurites. In addition, our data show that miR-183C regulates TG SN neurite growth, CX3CL1 and sP production, and the response to PA infection by directly targeting genes encoding these chemokine and neuropeptide and/or their processing enzymes as well as key receptor molecules mediating PA-induced activation.

## Methods

### Mice

All experiments and procedures involving animals and their care were reviewed and approved by the Wayne State University Institutional Animal Care and Use Committee and carried out in accordance with National Institute of Health and Association for Research in Vision and Ophthalmology (ARVO) guidelines (Approved protocol number: IACUC-22-05-4618). The study is reported in accordance with the Animal Research: Reporting of In Vivo Experiments (ARRIVE) guidelines. Euthanasia was performed by cervical dislocation under anesthesia with isoflurane followed by thoracotomy.

The miR-183C conventional KO – the miR-183C^GT/GT^ mice, are on a 129S2/C57BL/6-mixed background^49^ and were originally derived from a gene-trap (GT) embryonic stem cell clone from the German Gene Trap Consortium^49, 53, 54^.

The sensory nerve-specific conditional knockout, SNS-miR-183C-CKO mice [Na_v_1.8-Cre(+/-);miR-183C^f/f^;R26^LSL-RFP(+/+)^;Csf1r-EGFP(+/+)] and their WT littermates [Na_v_1.8-Cre(+/-);miR-183C^+/+^;R26^LSL-RFP(+/+)^;Csf1r-EGFP(+/+)] were produced by breeding of Na_v_1.8-Cre (+/-), miR-183C^f/+^, R26^LSL-RFP^ and Csf1r-EGF(+/-) mice as we described previously^27^. All breeding showed a normal mendelian inheritance pattern. The Na_v_1.8-Cre mice^55^ were kindly provided by Dr. John N. Wood, University College London, through Dr. Theodore J. Price, University of Texas, Dallas. This strain is now available at the Jackson Laboratory (Stock Number: 036564). The voltage-gated sodium channel Na_v_1.8 (encoded by the Scn10a gene) is one of the signature genes of the majority of nociceptive sensory neurons in the TG and dorsal root ganglia (DRG)^21, 56–58^. Na_v_1.8 promoter-driven Cre recombinase (Na_v_1.8-Cre) is expressed in nearly all corneal nociceptive sensory nerves^21, 55, 59^. Mice with miR-183C CKO allele, the miR-183C^f^, were provided by Dr. Patrick Ernfors, Karolinska Institutet, Sweden through the European Mouse Mutant Archive (EMMA. ID: EM12387). The miR183C^f^ allele has two loxP sites flanking the 5’ and 3’ ends of the miR-183C for robust Cre-mediated miR-183C CKO^60^.

The reporter strain, R26^LSL-RFP(+/+)^ mice^61^, also known as Ai14 (Stock number: 007914, the Jackson Laboratory) has a loxP-flanked STOP cassette (LSL) in front of a tdTomato red fluorescent protein (RFP) cassette, all of which are inserted into the ROSA26 locus^61^. The LSL prevents the transcription of tdTomato RFP; however, when Cre recombinase is present, the LSL cassette will be excised to allow the expression of tdTomato RFP. The Csf1r-EGFP, also called MacGreen mice^62^ were purchased from the Jackson lab (Stock number. 018549). In this strain, the EGFP transgene is under the control of the 7.2-kb mouse colony stimulating factor 1 receptor (Csf1r) promoter, allowing specific expression of EGFP in the mononuclear phagocyte system (MPS) myeloid cells, including monocytes (MCs), Mj and dendritic cells (DCs)^62, 63^. The Nav1.8-Cre-driven RFP and Csf1r-promoter driven EGFP in the SNS-miR-183C-CKO mice and their WT littermates ensure simultaneous double-tracing of sensory neurons and neurites (RFP+) and corneal resident myeloid cells (CRMCs) as we described previously^27^.

Sex is considered as a biological variable in all studies. 8-20 weeks old, male or female mice were used as separate groups in all experiments. n=5-6 mice/sex/genotype/experiment were used.

### Primary TG sensory neuron isolation

Primary sensory neurons were isolated from mouse TG using the 5-step OptiPrep gradient protocol as described before^64, 65^ with modifications. Briefly, after euthanasia, TG were carefully excised under a dissecting scope (VWR International, Radnor, PA) in a sterile biosafety hood (Microzone Cooperation, serial #BK-2-6). Two TG from a single mouse were subjected to enzymatic digestion in 750 μl of enzymatic digestion solution [EDS, 1 mg/ml Collagenase A (Cat 10 103 578 001. Sigma) and 2.4 U/ml Dispase II (Cat 17105041. Thermo Fisher Scientific. TFS) in Neurobasal A Media (NBM, Cat 12349015. TFS)] at 37 °C for 35 minutes (min) on a DisruptorGenie (Scientific Industries). The enzymes were deactivated by mixing with 750 μl of heat-inactivated fetal bovine serum (HI-FBS. HiClone. Cat No. SV30014-03) and mechanically dissociated to single-cell suspension by trituration using a P1000 pipet and a small-bore hole glass pipet. The single-cell suspension (in 1 ml in NBM) was layered on a 5-step OptiPrep gradient (Sigma-Aldrich. Cat No. D1556) and centrifuged for 30 min at 800 *g*. The lower end of the centrifuged gradient containing sensory neurons was then transferred to a new tube and washed twice with Wash Media [NBM + 2% B27 supplement (Invitrogen. Cat No. 10889-038) and 1% penicillin/streptomycin/neomycin (P/S/N. Invitrogen, Cat No. 15640055)]. The neurons were plated in poly-D-lysine (TFS, Cat No. A3890401)/laminin (TFS, Cat No. 23017015)-coated 24-well plates or the soma-chamber of custom-made microfluidic chambers (see below) in Plating Media [NBM + 2% B27 + 500 μM L-glutamine (GlutaMAX 1:100, TFS, Cat No. A12860-01), + 50 ng NGF (TFS, Cat No. 13290010) and 50 ng/ml GDNF + 1% P/S/N + 7.2 μM aphidicolin (Sigma. Cat No. A0781) + 40 μM fluorodeoxyuridine (Sigma. Cat No. F0503)]. After overnight culture, the plating media was replaced with Feeding Media [Plating Media without mitotic inhibitors aphidicolin and fluorodeoxyuridine] at 37 °C in 5% CO2 incubator for appropriate times before various treatments specified below. 2 hours before treatment, the Feeding Media was replaced with Treatment Media (NBM + 50 ng/ml NGF + 50 ng/ml GDNF).

### Microfluidic chambers

The custom-made microfluidic chambers (MFC) were manufactured as described previously^64^. Each MFC has two compartments (4x4x4 mm^2^), which we designated as soma and axon chambers, connected by an array of 100 micro-channels (10 μm width; 750 μm length; 2-2.5 μm height) (**Fig.1A**). Because of the restrictive height of the channels, cells plated in the soma chamber are unable to migrate through the channels. Standard soft lithography was performed using Sylgard 184 polydimethylsiloxane (PDMS) (Dow Silicones Corporation). PDMS microfluidics devices and glass-bottomed petri dish (MatTek corporation. Cat No. P50G-1.5-30-F) were exposed to plasma for 2 min at a flow rate of 5 cc/min under 200 mTorr vacuum pressure in a PE25-JW Benchtop Plasma Cleaning/Etching System (Plasma Etch). The plasma-exposed surfaces were immediately brought together to permanently bond the PDMS microfluidic device to the glass. The chambers were coated with poly-D lysine (PDL, 4 µg/cm².

**Figure 1.**
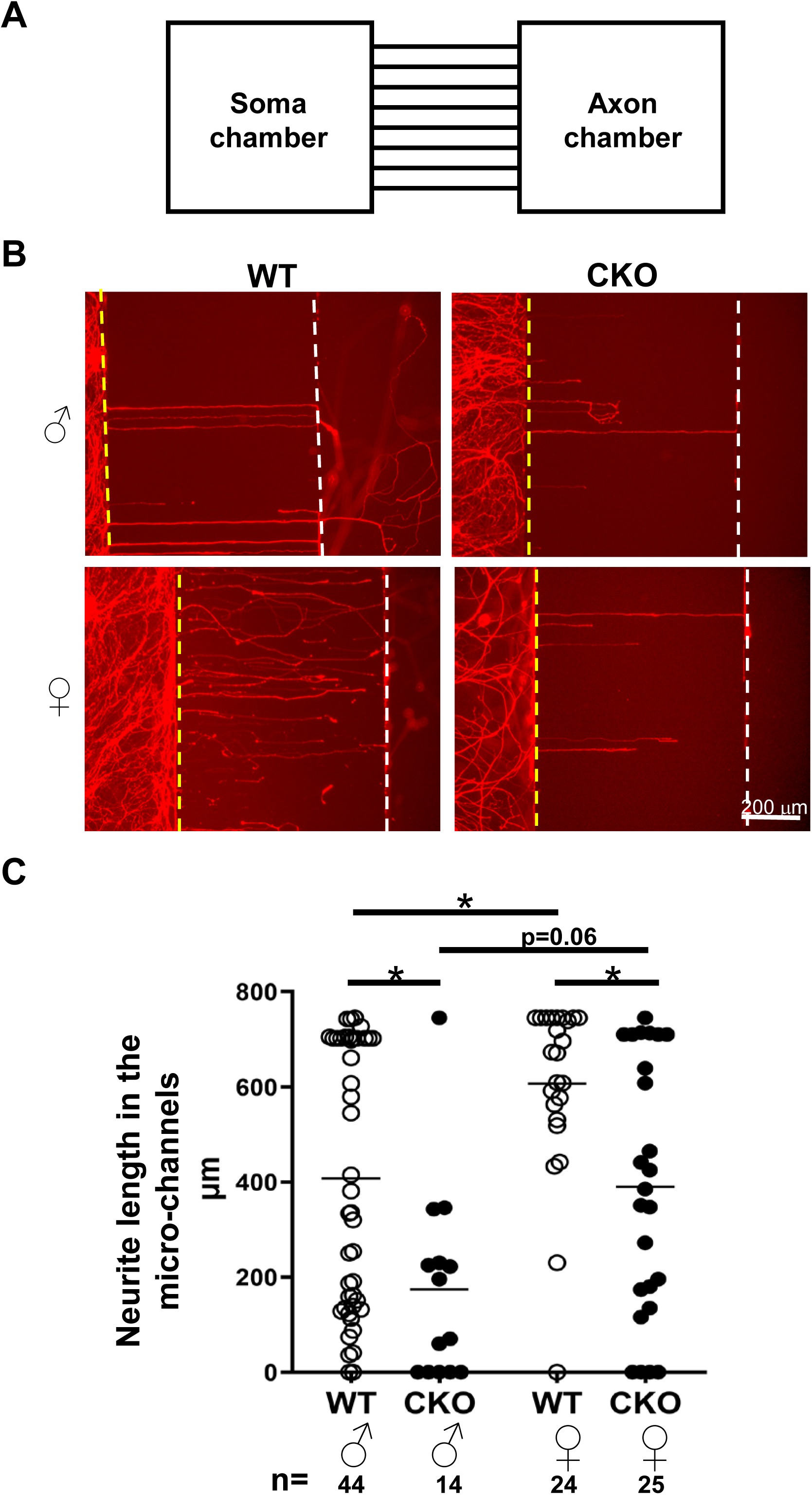
Neurite growth is decreased in the TG sensory neurons of SNS-miR-183C-CKO vs WT control mice. Neurite growth is increased in female vs male mice. A. Illustration of a custom-made microfluidic chamber. B. Representative photographs of neurites of TG sensory neurons of SNS-CKO (CKO) or their wild type control (WT) mice in microfluidic chambers. Dotted lines mark the boundaries of soma chamber (yellow) and the axon chamber (white). C. Statistical analysis of the neurite length in the microfluidic chambers. The bars in the graph mark the average. The numbers of neurites in the micro-channels of the MFCs (n) were labeled at the bottom of the figure.

TFS, Cat. No. A3890401) overnight and laminin (3 µg/cm². TFS, Cat. No. 23017015) for 1 hour, before the addition of cells and/or media.

### Neurite growth assay

For neurite growth assay, SNS-miR-183C CKO mice and their WT littermate controls, in which sensory nerves are labeled with Na_v_1.8-Cre driven RFP^27^, were used to isolate TG sensory neurons as described above. We first performed the neurite growth assay in the MFCs. Dissociated sensory neurons from each TG were plated in the soma chamber. To help guide the axons growing into the axon chambers through the micro-channels, the concentrations of NGF in axon chamber (500 ng/ml) was kept 10 times the ones in the soma chamber (50 ng/ml). 7 days later, the status of the neurite growth into the micro-channels was studied on an inverted fluorescent microscope (Leica). The length of all neurites in the micro-channels was measured manually in the photographs based on the scale bars. When no neurites grew into the micro-channels, the length of neurite growth was recorded as zero.

Second, we studied neurite length and growth in 24-well plates. To avoid overlapping of the neurites of individual neurons, low density culture (∼300 cells/well) was conducted in PDL and laminin-coated 24-well plate. Two and 4 days post-culture (dpc), neurons were studied and imaged using an inverted fluorescent microscope (Leica). Neurite tracing and quantification were performed using Fiji (ImageJ) with the NeuronJ plugin following the manufacturer’s instruction and as described previously^66, 67^. Neurites from at least 50 neurons/time-point/genotype/sex were quantified. Briefly, images were converted to 8-bit grayscale and spatial calibration was performed using a known scale bar. Individual RFP-labeled neurons were selected for analysis. Tracings were manually generated from the soma along primary neurites and branches. The *Measure Tracings* function was used to quantify total neurite length and number of branches.

### PA and LPS treatment of TG sensory neurons

TG sensory neurons were isolated from conventional miR-183C KO (miR-183C^GT/GT^) and their age- and sex-matched WT controls (n=5-6/genotype) as described above. Sensory neurons of two TG from each mouse were evenly distributed into 5-6 soma chambers of the MFCs as described above. 7 dpc, when the axons of TG sensory neuron had extended to the axon chambers, the axon chambers were subjected to one of the following conditions: 1) no-treatment control (NTC); 2) LPS (100 ng/mL, derived from PA-10. Sigma-Aldrich, Cat No. L8643); 3) PA [ATCC19660. 1.5x10^7^ colony forming units (CFUs)]; 4) TLR4 antagonist (20 µg/mL. LPS-RS Ultrapure. InvivoGen, Cat No. tlrl-prslps) for two hours before adding PA; 5) Formyl peptide receptor (FPR)1 antagonist (2 µg/mL. Boc-MLF. TOCRIS, Cat No. 67247-12-5) for two hours before adding PA; 6) Antagonists of both TLR4 and FPR1 for two hours before adding PA. After overnight (∼16 hours) treatment, the supernatant media of both axon and soma chambers and the cell lysate of the soma chambers were harvested for ELISA assays of sP [R&D System, Cat No. KEG007; minimum detectable dose (MDD) 16.8-43.8 pg/mL] or CX3CL1 (R&D Systems, Cat No. MCX310. MDD 0.08–0.32 ng/mL) as we described previously^27^.

### Target luciferase reporter assay

Target luciferase reporter constructs of mouse Tac1 (Cat No. MmiT0965624-MT06), Cx3cl1 (Cat No. MmiT028162-MT06), Adam10 (Cat No. MmiT090821-MT06), Tlr4 (Cat No. MmiT088511a/b-MT06), Nrp1 (Cat No. MmiT093902-MT06), and Fpr1 (Cat No. HmiT149043-MT06), which carry the 3’ UTR of these genes with predicted target sites for miR-183/96/182 downstream of a firefly luciferase (FL) cassette in the pEZX-MT06 vector, were purchased from Genecopoeia. The pEZX-MT06 vector also carries a constitutively expressed Renilla luciferase (RL) cassette for transfection control (Genecopoeia). Target luciferase reporter assays were performed as described previously^51^. Briefly, 50 ng of the construct plasmid DNA were co-transfected with miRNA mimics [mmu-miR-183, or -182, or -96, or miR-183/96/182, or negative-control oligo duplex with scrambled sequences (scr mimic)] (10 nM; Ambion/Thermofisher Scientific) or miRCURY LNA miRNA Power Inhibitors [anti-mmu-miR-183, -182, -96 or anti-mmu-miR-183/96/182 or negative control oligonucleotides with scrambled sequences (scr anti-miR)] (10 nM; Exiqon/Qiagen)] into the 50B11 rat dorsal root ganglion cell line^68^ (kindly provided by Dr. Ahmet Hoke, the Johns Hopkins University) using Lipofectamine 2000 (Invitrogen) as we described previously^69^. Forty-eight hours later, we harvest the cells for dual luciferase assays using the Luc-Pair miR Luciferase Assay System (Genecopoeia) and a Glomax plate reader (Promega, Madison, WI, USA). Relative luciferase activity was calculated as FL activity normalized by RL activity. All experiments were performed with 5 replicates under each condition (n=5).

### Statistical analysis

Statistical analyses were performed as described previously^27, 51^. When the comparison was made among more than 2 conditions, ordinary one-way ANOVA with Bonferroni post hoc test was employed *(GraphPad Prism)*; adjusted p<0.05 was considered significant. Otherwise, a two-tailed Student’s *t* test was used to determine the significance; p<0.05 was considered significant. Each experiment was repeated at least once to ensure reproducibility and data from a representative experiment are shown. Quantitative data is expressed as the mean ± SEM.

## Results

### Inactivation of miR-183C decreases neurite length/growth in culture

To characterize the impact of miR-183C inactivation in sensory neurons on their neurite growth, we harvested TG sensory neurons from young adult SNS-miR-183C CKO mice and their WT control littermates, in which sensory nerves are labeled with TdTomato RFP. We first performed a neurite growth assay in the MFC chambers (**Fig.1A**), in which TG sensory neurons were plated in the soma chamber and 7 days later, we measure the neurite lengths in the micro-channels. Our results showed that the average neurite length in the micro-channels of TG neurons from SNS-CKO vs WT controls was significantly decreased in both male and female mice (**Fig.1B&C**): the average length of the neurites was decreased from 407.5 ± 41.6 μm (Mean ± SEM, n= 44) in WT to 174.1 ± 55.9 μm (n=14) CKO of male mice, and from 606.7 ± 37.7 μm in WT (n=24) to 389.84 ± 54.1 μm in CKO (n=25) of female mice (**Fig.1C**). When compared to male mice, the neurite length in the micro-channels of TG sensory neurons of female mice was significantly increased (**Fig.1C**), suggesting a sex-dependent difference in neurite growth.

To further evaluate the role of miR-183C in neurite growth, we plated TG sensory neurons in 24-well plates at low density to avoid neurite overlapping and performed neurite tracing at 2 and 4 dpc (**Fig.2A**). The experiment was first performed in male mice (**Fig.2A-C**). Our results showed that, at 2 dpc, the total neurite length per sensory neuron was significant decreased by ∼45.7% in the miR-183C-CKO vs their WT controls (from 4891.6 ± 330.6 μm in WT to 2656.9 ± 281.8 μm in SNS-CKO mice. n=53 of both WT and CKO mice. p<0.05) (**Fig.2B**). The number of branches per neuron was also decreased (by ∼ 44.6%) in the SNS-CKO vs WT controls (from 41 ± 3 in WT to 24 ± 6 in CKO) (**Fig.2C**). From 2 to 4 dpc, the total neurite length and the number of neurites per neuron were increased in both CKO and WT mice (**Fig.2B&C**).

**Figure 2.**
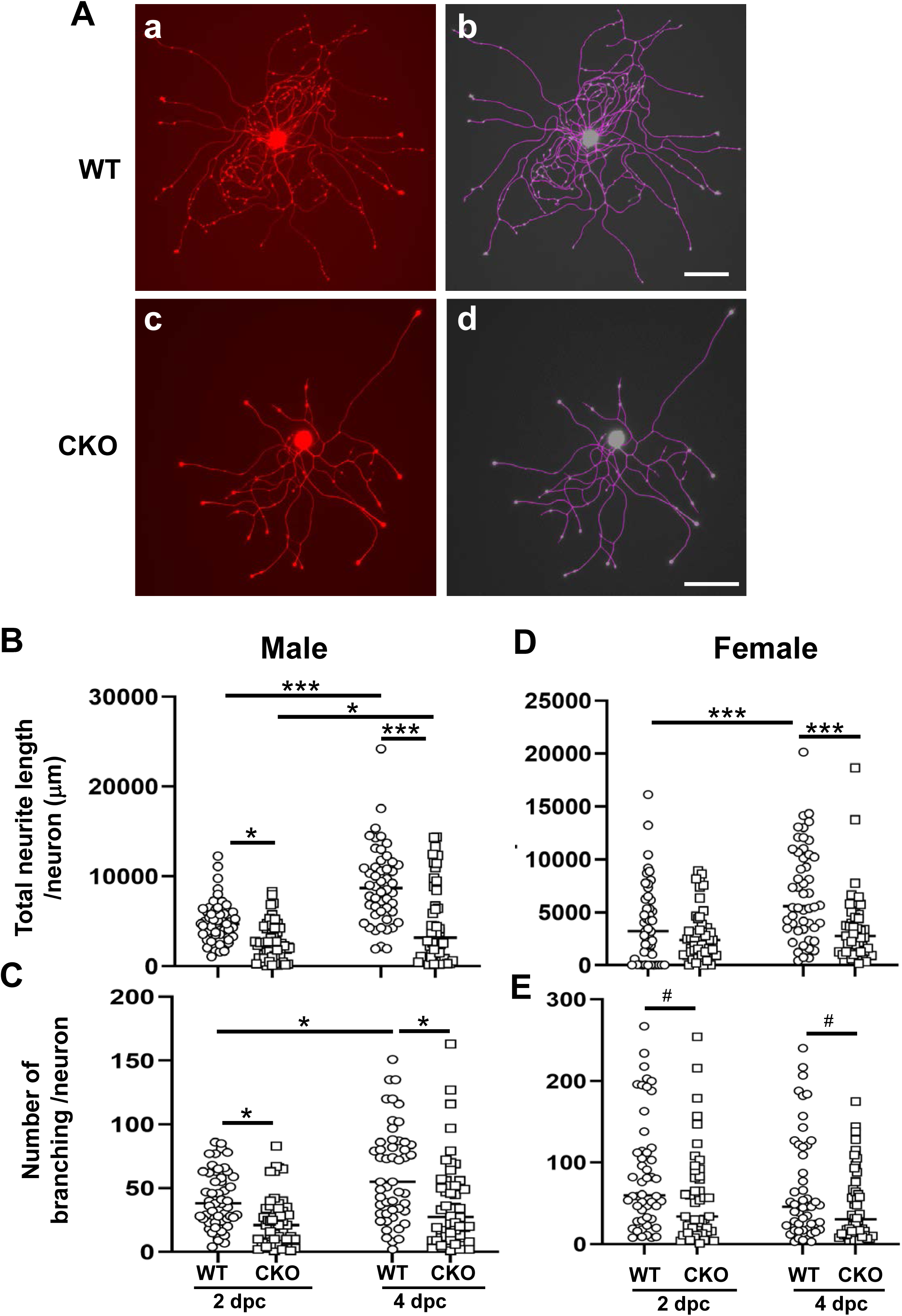
Inactivation of miR-183C resulted in decreased neurite length and number of branches of TG sensory neurons. **A.** Representative microscopic photographs (a,c) and neurite tracing using the Neuroanatomy-Fiji ImageJ software (b,d) of TG sensory neurons of male WT (a,b) and SNS-CKO mice after 2 days post-culture (dpc). Scale bars: 200 μm. **B-E**. Quantification of total neurite length per neuron (B&D) and number of branches per neuron (C&E) of TG sensory neurons of male (B&C) or female (D&E) mice at 2 and 4 dpc. n=53/genotype at 2 dpc; 50/genotype at 4 dpc. *: p<0.05, ***:p<0.001 by one-way ANOVA of multiple sample comparison with Bonferroni post hoc test*. #:* p<0.05 by Student’s *t* test.

However, the increment of neurite length was smaller in CKO vs WT mice. The neurite length was increased by ∼75.2% to 4794.1 ± 631.0 μm (p<0.05) in CKO mice, while ∼85.1% to 8996.0 ±609.2 μm (p<0.0001) in WT mice (**Fig.2B**). At 4 dpc, comparing the SNS-CKO vs WT mice, the total neurite length per neuron continued to be decreased in SNS-CKO vs WT controls (by ∼46.7%. p<0.001), while the number of neurite branches was decreased by ∼39.7% (p<0.05). These data suggest that inactivation of miR-183C in TG sensory neurons results in decreased neurite length and branching.

Subsequently, we performed neurite tracing experiment in female SNS-CKO and WT control mice. Consistent with the results in males, similar trends were observed in neurite length and grow in female mice (**Fig.2D&E**): the total neurite length and number of branches was decreased in SNS-CKO vs WT mice.

In the second neurite tracing assay, to compare TG sensory neurons from male and female samples simultaneously, male samples were also collected and observed side-by-side with the female samples. Comparing male vs female samples, our result show that, consistent with the results by the MFC neurite growth assay (**Fig.1**), both the total neurite length and the number of neurite branches per neuron showed an increased trend in the female vs male mice (**Fig.S1**), suggesting sex-dependent difference in TG sensory neurons.

### Inactivation of miR-183C in TG sensory neurons enhances their response to PA infection

Previously, we showed that miR-183C directly targets key receptor molecules, e.g., TLR4, resulting in enhanced basal production of proinflammatory cytokines in corneal resident Mφ^51^.

Recently, we further reported that miR-183C also target Cx3cl1, a unique neuron-produced transmembrane chemokine, suggesting a role in the neuroimmune interplay in the cornea^27^. Additional target prediction analysis suggests Tac1, the gene encoding neuropeptide sP, is a predicted target gene of miR-183 and miR-96 (see below in Fig.5B & Fig.S3A). These observations support the hypothesis that miR-183C imposes direct regulation on neuropeptide and chemokine production and the response of TG sensory neurons to PA infection.

To directly test this, we performed PA and LPS treatment assays in the microfluidic chambers. To simulate PA infection of the cornea, the TG sensory neurons of miR-183C KO or age- and sex-matched WT littermate controls were plated in the soma chambers. 7 dpc, when the neurites had grown into the axon chambers, we infected the neurites in the axon chambers with PA (ATCC19660; 1.5x10^7^ CFU). To test whether lipopolysaccharide (LPS), a major component of the PA cell wall, mediates the PA-induced TG neuronal response, we also challenged the neurites with LPS (100 ng/ml) in parallel. After an overnight challenge (∼16 hours), we harvested the supernatants of the axon chambers and the TG neuronal cell lysates and performed ELISA assay for CX3CL1 and sP. Our results showed that, under NTC condition, the basal levels of CX3CL1 were increased in both the supernatant of the axon chamber (2.60±0.35 ng/ml) and the lysate of sensory neuronal cell bodies of miR-183C KO (1.63±0.11 ng/ml) vs WT control mice (0.42±0.06 ng/ml, and 0.54±0.14 ng/ml, respectively) (**Fig.3A&B**), consistent with the hypothesis that Cx3cl1 is a direct target of the miR-183C^27^. Although LPS alone did not cause a significant induction of the CX3CL1 in WT mice, it significantly increased the levels of CX3CL1 in both the supernatant of the axon chamber and the cell lysate in the soma chamber of the KO mice (**Fig.3A&B**). These data suggest that the miR-183C regulates CX3CL1 production in TG sensory neurons and their response to LPS stimulation; and that LPS has a modest role in mediating the PA-induced TG sensory neuronal response of CX3CL1 production/secretion.

**Figure 3.**
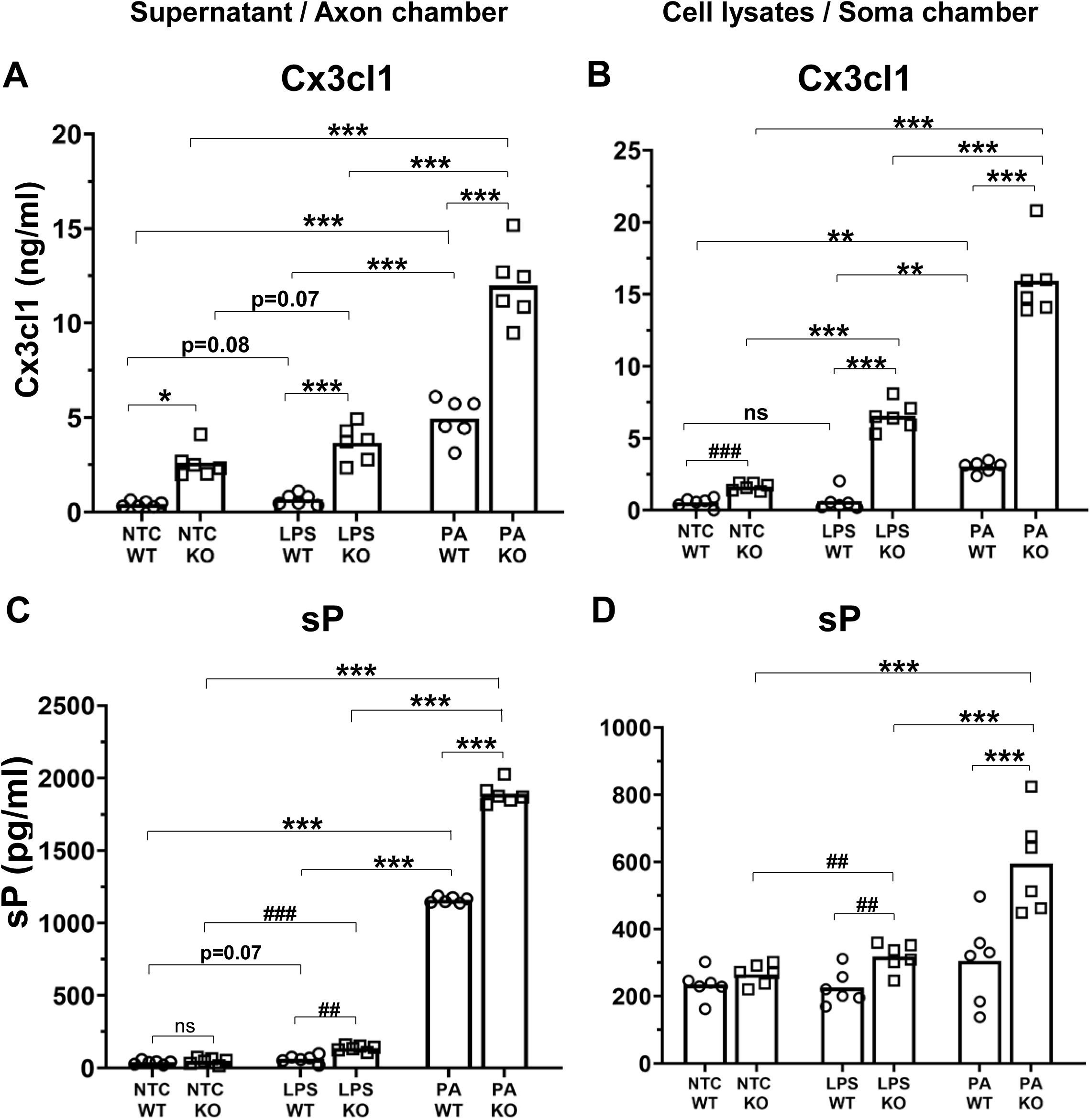
PA infection of TG neurites and nerve endings induced the expression of neuropeptides, CX3CL1 (A) and sP (B); and inactivation of miR-183C enhanced basal level and PA-induced production of CX3CL1 and sP. ELISA assays in the supernatant of the axon chambers of the axon chamber (A&C) and the cell lysate of the soma chamber (B&D). NTC: no treatment control. *: p<0.05, ***:p<0.001 by one-way ANOVA of multiple sample comparison with Bonferroni post hoc test. ns: not significant; ##: p<0.01; ###: p<0.001 by Student’s *t* test

In contrast to LPS treatment, PA bacteria induced significant upregulation of CX3CL1 in both WT and KO sensory neurons and in both the axon-chamber supernatant and the cell lysates of the soma chamber, in a higher amount than LPS alone, with the levels of CX3CL1 in the axon chamber of KO mice (12.0±0.9 ng/ml) ∼ 2.4x folds higher than that of WT mice (4.9±0.5 ng/ml) (**Fig.3A&B**). These data suggest that, first, PA bacteria directly activated TG sensory neurons and induced their production/secretion of CX3CL1; second, other component(s) of PA, in addition to LPS, mediate the PA-induced TG sensory neuronal response; third, miR-183C regulates CX3CL1 production and secretion by TG sensory neurons.

For sP, its expression level did not show significant upregulation in either the supernatant of the axon chambers or the cell lysate of the soma chamber of the NTC control samples of KO vs WT mice. Similar to CX3CL1, although LPS alone did not have significant induction of sP in WT TG sensory neurons, it showed moderate enhancement of sP levels in both the axon-chamber supernatant and the soma-chamber lysates of the KO mice, resulting in increased levels of sP in KO vs WT mice in response to LPS treatment (**Fig.3C&D**). PA infection resulted in a higher induction of sP production in the both WT and KO mice, in both the axon-chamber supernatant and the soma-chamber lysates, with the levels of sP significantly increased in the KO vs WT mice (**Fig.3C&D**). Similar to its role in Cx3cl1 production, these data suggest that miR-183C regulates sP production in the TG sensory neurons, and that exposure of TG sensory neurites to PA bacteria induces sP production/secretion; LPS alone has a modest role in mediating this response.

### TLR4 and Fpr1 mediate TG sensory response to PA infection

Previous reports suggested that bacteria can activate sensory neurons through specific receptors^70–78^, including TLR4, and formyl peptide receptor (Fpr1)^72, 73^. We showed that TLR4 is a direct target of miR-183C in corneal resident Mφ^51^. Additional target prediction suggest that Fpr1 is also a potential direct target of miR-183C (See below in Fig.5F & Fig.S3C). Therefore, we hypothesized that TLR4 and FPR1 may mediate PA-induced increased production of CX3CL1 and sP. To test this hypothesis, we pre-treated the neurites in the axon chambers with a TLR4-specific antagonist, TLR4-RS (InVivoGen), and/or a FPR1-specific antagonist, Boc-MLF (TOCRIS) 2 hours before PA infection. Our results showed that TLR4-RS or Boc-MLF alone significantly inhibited PA-induced enhanced production and secretion of CX3CL1 and sP in the axon-chamber supernatant (**Fig.4**) and the cell lysate in the soma chamber (**Fig.S2**). Comparing the effects of TLR4-RS with the ones of Boc-MLF, the latter showed a higher inhibition on PA-induced production and secretion of both CX3CL1 and sP in the axon-chamber supernatant (**Fig.4**). Pre-treatment of the TG sensory neurites simultaneously with both TLR4-RS and Boc-MLF in the axon chamber resulted in complete inhibition of PA-induced enhanced production and secretion of CX3CL1 and sP (**Fig.4**), suggesting that TLR4 and FPR1 together are fully responsible for PA-induced CX3CL1 and sP production and secretion. For CX3CL1, simultaneous treatment of both TLR4-RS and Boc-MLF resulted in further reduction of its levels in the supernatant of the axon chamber of both WT (0.337±0.016 ng/ml) and KO mice (0.563±0.027 ng/ml) comparing to their corresponding NTC control samples (0.635±0.066 ng/ml in WT; 1.079±0.047 ng/ml in KO mice) (**Fig.4A**), suggesting that TLR4 and FPR1 also modulate the basal production and secretion of CX3CL1 in TG sensory neurons at homeostatic condition.

**Figure 4.**
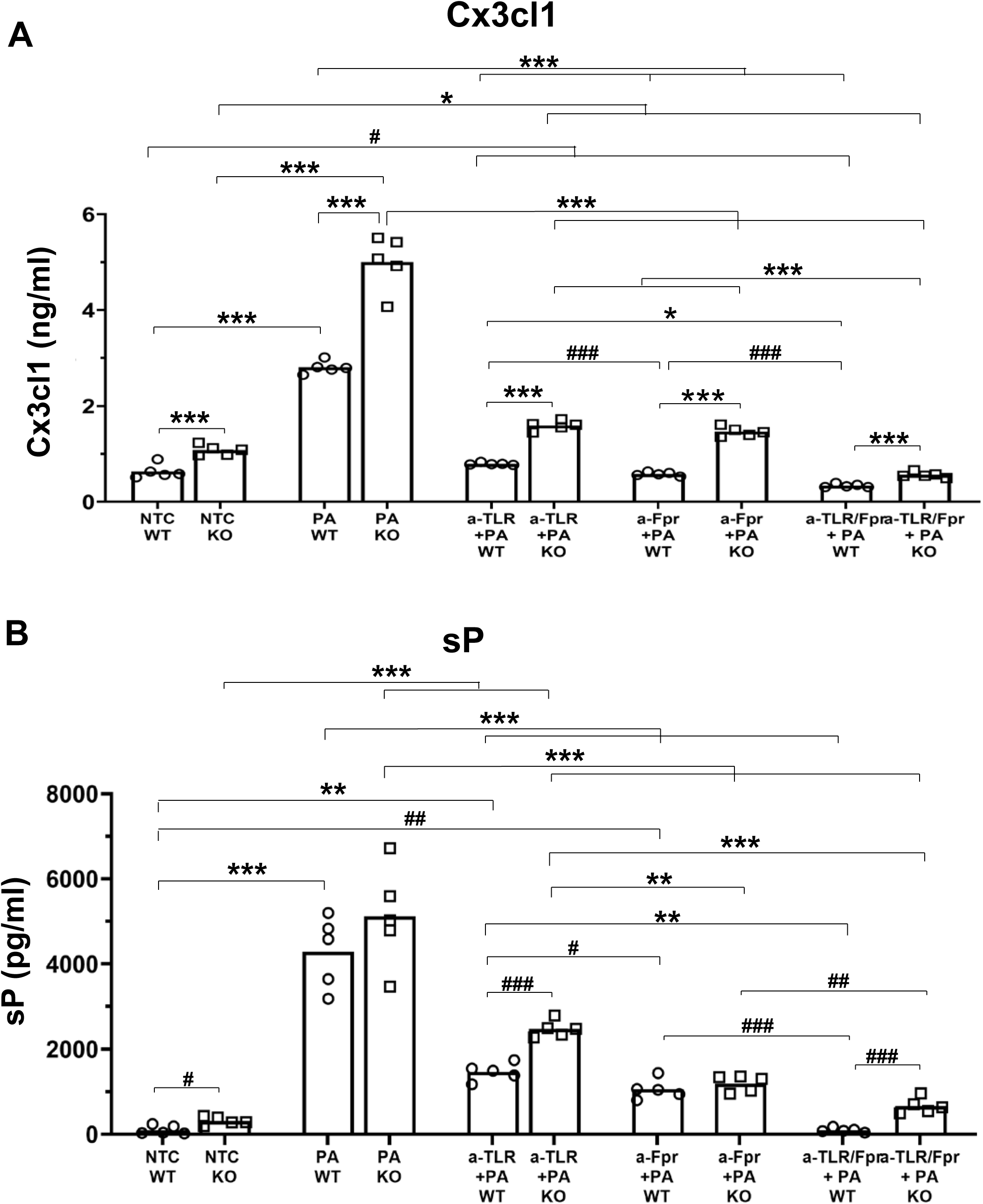
TLR4 and FPR1 mediate PA-induced expression of neuropeptides, CX3CL1 and sP. ELISA assays in the supernatant of the axon chambers. NTC: no-treatment control; a-TLR+PA: the axon chamber pretreated with TLR4-specific antagonist, LPS-RS Ultrapure (20 µg/mL) 2 hours before PA infection; a-Fpr+PA: the axon chamber pre-treated with Fpr1 antagonist, Boc-MLF (2 µg/mL) 2 hours before PA infection; a-TLR/Fpr+PA: the axon chamber pre-treated with both LPS-RS Ultrapure (20 µg/mL) and Fpr1 antagonist, Boc-MLF (2 µg/mL) 2 hours before PA infection. *: p<0.05, **: p<0.01, ***:p<0.001 by one-way ANOVA of multiple sample comparison with Bonferroni post hoc test. #: p<0.05, ##: p<0.01, ###: p<0.001 by Student’s *t* test.

### miR-183C targets key genes involved in neurite growth, neuropeptide production and bacterium-induced sensory neuronal activation

Previously, we showed that Neuropilin (Nrp)1^79^, a negative regulator of CSN growth into the cornea during development^80, 81^ and CSN regeneration in epithelial wound healing^82, 83^, is a predicted target of miR-183 and miR-182^52^. Consistently, Nrp1 is upregulated in the TG of naïve miR-183C ko mice^52^, suggesting that miR-183C targets Nrp1 in TG neurons, which may contribute to the miR-183C KO- and knockdown-induced reduction of CSN density that we reported previously^28, 52^ and decreased neurite length/growth demonstrated in **Fig.1&2**. To validate that miR-183C directly targets Nrp1, we performed target luciferase reporter assay using a construct carrying mouse Nrp1 3’UTR sequence in a dorsal root ganglion sensory neuronal cell line, the 50B11 cells^68^. Our result showed that over-expression of miR-182 and miR-183 specifically suppressed the expression, while knockout of miR-182/183 as well as miR-96 disinhibited the expression of Nrp1 (**Fig.5A**), confirming that Nrp1 is a direct target of miR-183C.

**Figure 5.**
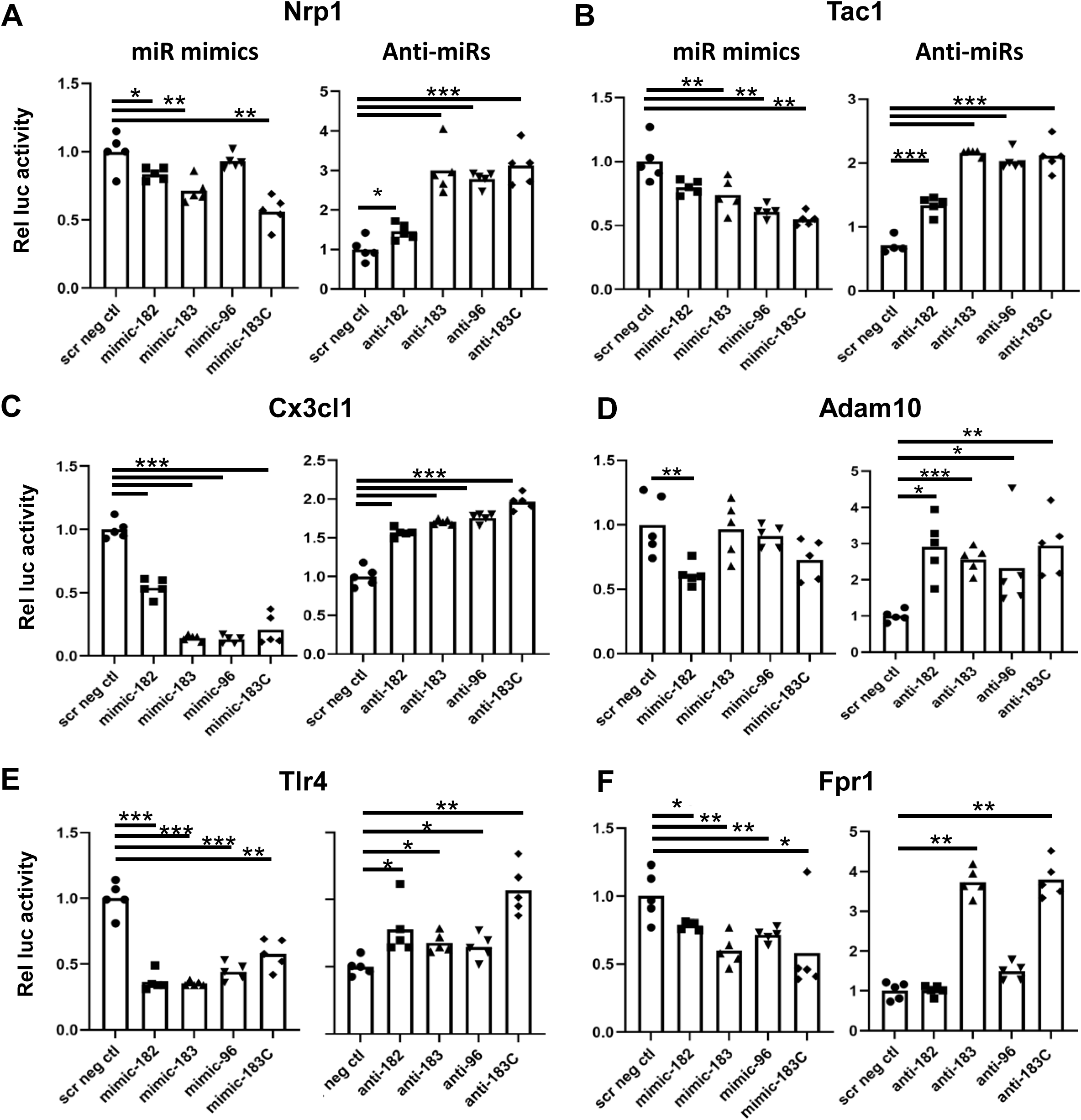
miR-183C target Nrp1, Tac1, Cx3cl1, Adam10, Tlr4, and Fpr1. Target luciferase assays of mouse Nrp1 (A), Tac1 (B), Cx3cl1 (C), Adam10 (D), Tlr4 (E), and Fpr1 (F). *: p<0.05; **: p<0.01; ***: p<0.001.

Additional target prediction using the TargetScan algorithm^84–88^ suggested that Tac1, the precursor gene for sP, is a predicted target gene of two members of the miR-183C, miR-183 and miR-96 (**Fig.S3A**). The increased expression of sP by the TG sensory neurons of miR-183C KO vs WT mice upon LPS treatment and PA infection (**Fig.3B**) supports the hypothesis that Tac1 is targeted by miR-183C in TG sensory neurons. Consistent with this hypothesis, our target luciferase reporter assays showed that over-expressing of miR-183, miR-96 or miR-183C by miRNA mimic significantly suppressed, while knockdown of all three miRNAs by anti-miRs significantly dis-inhibited the expression of Tac1 (**Fig.5B**), validating that Tac1 is a direct target of miR-183C.

Previously, we reported that Cx3cl1 is a predicted target of miR-183^27^. To validate this, we performed target luciferase reporter assay and showed that over-expression of miR-183, as well as miR-182 and miR-96, inhibited Cx3cl1 expression; while knockdown of miR-183, 182, and/or 96 disinhibited Cx3cl1 expression (**Fig.5B**), confirming that miR-183C directly regulates that expression of Cx3cl1 at a post-transcriptional level. The fact that miR-182 and miR-96 also targeted Cx3cl1 in this assay is possibly a result of the overlapping functions of the three miRNAs in the miR-183C because of high sequence homology^48, 49^.

Cx3cl1 is known as an unusual member of the chemokine family. It is synthesized as a type I transmembrane protein, with a long ectodomain containing the CX3C chemokine domain displayed on an extended, highly glycosylated, mucin-like stalk^89–91^. Proteolytic cleavage by the transmembrane proteases of ADAM (a disintegrin and metalloproteinase)-family ADAM10 and ADAM17 results in shedding of the ectodomain forming the soluble form of CX3CL1 (sCX3CL1) ^92, 93^. ADAM10 is mostly responsible for the constitutive shedding of sCX3CL1^92^. Additional target prediction suggests that ADAM10 is a direct target of miR-182 (**Fig.S3B**). Consistently, Adam10 is significantly upregulated in the TG of both miR-183C KO and the SNS-CKO mice when compared to their corresponding WT controls (data not shown). To further validate the miRNA-target relationship, we performed target luciferase reporter assay and confirmed that Adam10 is targeted by miR-183C (**Fig.5D**), suggesting miR-183C is involved in the processing/release of sCX3Cl1, at least in part, by targeting Adam10.

Previously, we showed that Tlr4 is predicted to be targeted by all three miRNAs of the miR-183C in corneal resident Mφ^51^. Additional target prediction suggest that Fpr1 is also a potential direct target of miR-183 (**Fig.S3C**). To test these predictions, we performed target luciferase assays of Tlr4 and showed that miR-182, -183 and 96 mimics robustly inhibited, while anti-miRs of all three miRNAs un-inhibited the luciferase reporter activities. For Fpr1, mimics of all three miRNAs inhibited the luciferase activities, with the most inhibition by miR-183; consistently, anti-miR-183 specifically disinhibited the luciferase reporter activity, consistent with the prediction that Fpr1 is targeted by miR-183. These data validated that miR-183C regulates the key receptor molecules mediating PA-induced TG sensory neuronal response.

## Conclusion and Discussion

Increasing evidence suggests that corneal sensory innervation modulates the corneal immune response to PA infection by releasing neuropeptides^21–26^; however, whether and how CSN are activated directly by PA bacteria and involved in the initiation of PA keratitis is still unknown.

Here in this study, we provide experimental evidence that miR-183C regulates TG sensory neurite growth; PA directly interact with TG sensory neurites/nerve endings and activate the production and secretion of neuron-produced chemokine CX3CL1 and neuropeptide sP, both of which are key to modulating the corneal niche and neuroimmune interactions in both homeostatic and pathological conditions^22, 23, 25, 29, 94, 95^; and that miR-183C regulates the basal as well as PA-induced production and secretion of these key chemokine and neuropeptide by directly targeting genes involved in these processes. To our knowledge, this is the first report demonstrating that PA directly activate TG sensory neurons through their neurite and/or nerve endings and inducing their production and secretion/release of CX3CL1 and sP. In 2021, Lin et al. reported that PA activated TG neurons by calcium imaging and enhanced secretion of CGRP^21^; however, in this report, PA was applied directly to the TG neuronal culture with both soma and neurites, which did not distinguish whether the activation was achieved by PA interaction with neuronal cell bodies or neurites/nerve endings, the latter of which simulates the activation of TG sensory neuron by PA infection in the cornea. Furthermore, this report was retracted in 2024^96^.

First, using custom-made MFCs^64^, we showed that the average neurite length projecting into the microchannels by TG sensory neurons of the SNS-miR-183C-CKO mice is significantly decreased when compared to the WT controls in both male and female mice. This result is corroborated by the findings of the neurite tracing experiment, suggesting that miR-183C regulates TG sensory neurite extension. In addition, the neurite tracing experiment also showed that the number of branches per neuron was also significantly decreased in TG sensory neurons of SNS-CKO vs WT controls, suggesting that miR-183C imposes dual regulation of both TG sensory neurite extension as well as branching. These data, to a degree, provide an explanation to the phenotypes that we reported early that inactivation of miR-183C results in decreased CSN density in the miR-183C conventional KO^28^ and the SNS-CKO mice^27^. In search of potential mechanism underlying miR-183C’s regulation on TG sensory neurite growth, we identified and validated that Nrp1, a crucial regulator of neurite growth, axon guidance and branching^79–83, 97^, is a direct target gene of miR-183C, suggesting that miR-183C’s regulation of corneal sensory innervation is, at least in part, mediated by Nrp1. However, we do not expect that Nrp1 is the sole mediator of this function, since our previous studies have shown that more than a dozen of genes involved in neuronal projection and axon guidance are targeted by miR-183C in mouse TG^27^. Therefore, we believe that miR-183C’s regulation on corneal sensory innervation is achieved by its simultaneous targeting and quantitative modulation of a collection of genes involved in neuron projection and axon guidance (**Fig.6**). Further studies will be required to fully test this hypothesis.

**Figure 6.**
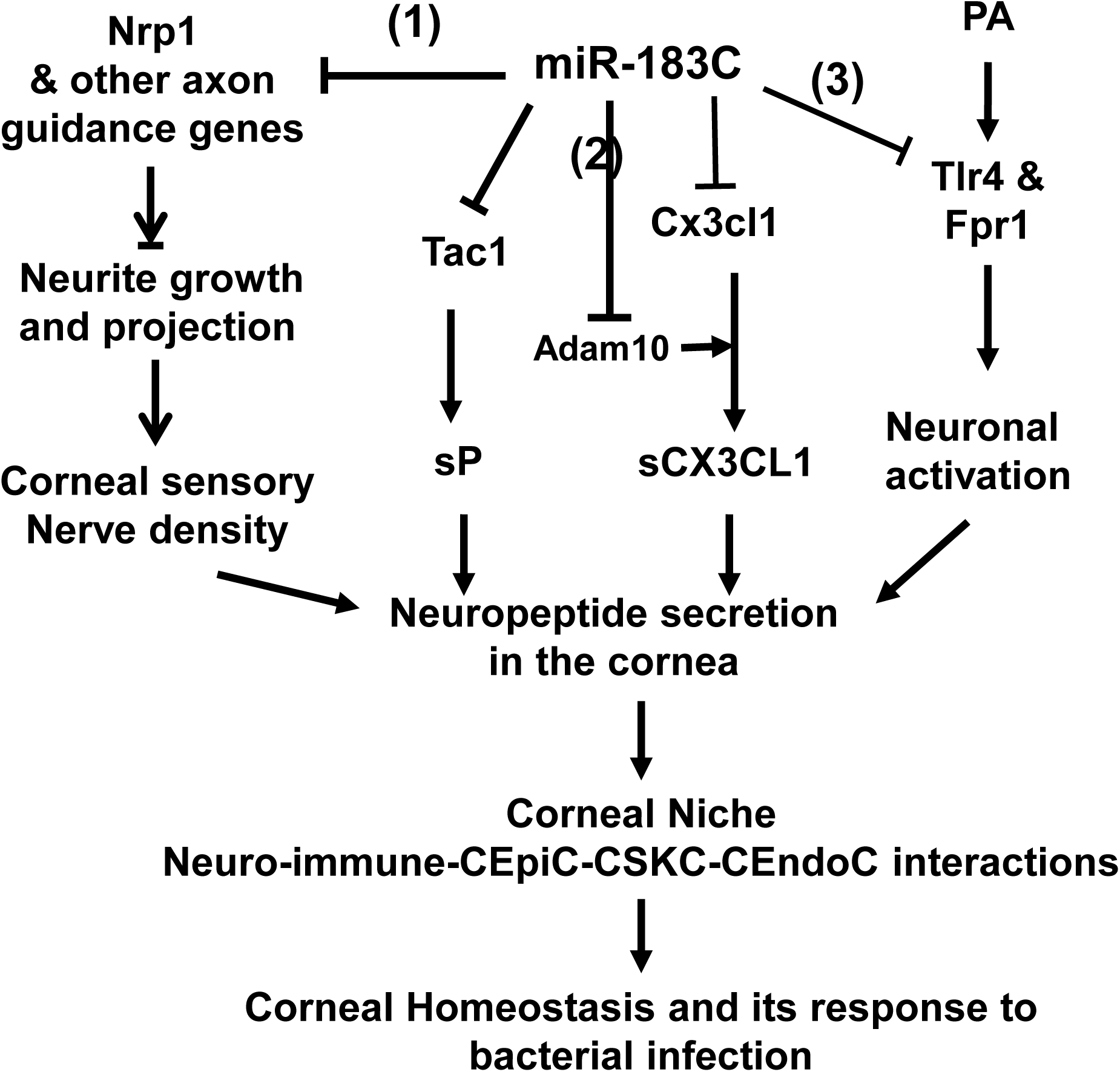
Illustration of molecular pathways by which miR-183C modulates corneal niches and its response to PA infection through its regulation of TG sensory neuronal functions. (1) miR-183C modulates corneal sensory nerve density by regulating TG sensory neuron neurite growth and projection through targeting Nrp1 and other neurite growth- and axon guidance-related genes. (2) miR-183C also regulates the expression of neuropeptides, e.g. sP and CX3CL, through directly targeting the genes encoding sP, Tac1, and Cx3cl1, and Adam10. In addition, (3) miR-183C modulates PA-induced TG sensory neuronal activation and/or neuropeptide releases by targeting receptor genes, e.g., Tlr4 and Fpr1. Through these pathways, miR-183C regulates the levels of neuropeptides in the cornea, modulating the corneal niche and interactions among all cellular components of the cornea, and therefore, corneal homeostasis and its response to bacterial infection. CEpiC: corneal epithelial cells; CSKC: corneal stromal keratocytes; CEndoC: corneal endothelial cells.

In addition to the differences between miR-183C KO and WT mice, we unexpectedly observed a sex-dependent difference that the total neurite length and number of branches of TG sensory neurons are increased in female vs mice. This result is consistent with a previous report in the mouse^98^ showing that CSN of female mice regenerate faster than male mice after corneal epithelial debridement^98^. Similar sex-dependent difference has also been reported in rat in various peripheral nerve crushing models^99–101^. It will be of importance to further validate these sex-dependent differences of sensory innervation.

Using the custom-made MFC^64^, we separated the neurites and nerve endings in the axon chamber from the cell bodies of the TG sensory neurons in the soma chamber. This allowed us stimulate the TG sensory neurons through their neurites/nerve endings in the axon chamber and study their response in the axon chamber – an in vitro system simulating TG sensory neuronal response to PA infection in the cornea. We showed that, in TG sensory neurons from WT mice, PA bacteria, but not LPS alone, significantly induced the production and secretion of key neuropeptides, CX3CL1 and sP, in both the soma lysate and the supernatant of the axon chamber. These results suggest that engagement of PA bacteria with sensory neurites and nerve endings directly activated sensory neurons to enhance their production (in soma lysate) and secretion or release of the neuropeptides (in axon-chamber supernatant). Our data suggest that LPS alone is not the major mediator of PA-induced neuropeptide response of the TG sensory neurons; other component(s) of PA bacteria are involved in this process.

Inspired by previous studies suggesting that TLR4 is involved in LPS-induced sensitization of TG sensory neurons^73^, while FPR1 mediates *Escherichia* (E.) *coli*-derived and *Staphylococcus* (S.) *aureus*-derived N-formylated peptides-induced activation of dorsal root ganglion sensory neurons^72^, we tested whether TLR4 and FPR1 mediate PA-induced neuropeptide response of TG sensory neurons. We pre-treated the neurite/nerve endings in the axon chamber of the TG sensory neurons with TLR4-specific and/or FPR1-specific antagonists before PA bacteria were added. Our results showed both TLR4- and FPR1-specific antagonists significantly reduced PA-induced neuropeptide production and secretion; FPR1-specific antagonist exhibited stronger inhibition of this response than TLR4-specific antagonist; and simultaneous application of both antagonists completely inhibited PA-induced neuropeptide production and secretion. These data suggest that both TLR4 and FPR1 contribute to PA-induced CX3CL1 and sP production and secretion, with FPR1 playing a larger role. TLR4 and FPR1 together are fully responsible for mediating PA induced neuropeptide response of TG sensory neurons. Intriguingly, simultaneous treatment of both TLR4- and FPR1-specific antagonists resulted in further reduction of CX3CL1 level comparing to the NTC control samples, suggesting that TLR4 and FPR1 also modulate the basal production of CX3CL1 of TG sensory neurons at homeostatic condition.

In this study, we focus on these two neuropeptides because they are known to play important roles in neuroimmune interaction. The full-length transmembrane CX3CL1 plays important roles in cell-cell adhesion^102–104^, while sCX3CL1 acts as a chemotactic factor inducing chemotaxis^89, 90^. CX3CL1 is known to promote chemotactic migration of microglia in the central nervous system (CNS)^105, 106^, mediate the homing of resident myeloid cells in the cornea^29^ and recruitment of Mφ in various tissues and pathological conditions^107^. sP causes immune/inflammatory response by binding to neurokinin-1 receptor (NK1R) on immune cells, endothelial cells and corneal constituent cell types to enhance microvascular permeability, pro-inflammatory cytokine production, and monocyte and neutrophil infiltration^22, 23, 25, 94, 95, 108, 109^. Considering that the cornea contains tens of thousands of corneal sensory nerve endings embed between the surface epithelial cells^11, 14, 17–20^, we predict that direct contact of PA bacteria with these densely distributed sensory nerve endings will rapidly elevate the levels of these key pro-inflammatory neuropeptides, which initiates a chain of immune/inflammatory response in the cornea, leading to PA keratitis. Therefore, our data provide a potential molecular mechanism underlying sensory neurons’ roles in the initiation of the pathogenesis of PA keratitis.

Mechanistically, we identified that the gene encoding the precursor of sP, Tac 1, is a direct target gene of miR-183C. In addition, the genes encoding CX3CL1 itself as well as the transmembrane proteases metalloproteinase ADAM10, which is responsible for the constitutive shedding of sCx3cl1^92^, were both targeted by miR-183C. These data suggest that miR-183C regulates the basal expression of Cx3cl1 and sP by directly targeting the genes encoding these neuropeptides themselves and/or the enzyme(s) involved in their biogenesis (**Fig.6**). The increased basal level of sCX3CL1 in the axon chamber of TG sensory neurons of the KO vs WT mice is consistent with our previous report that CX3CL1 is increased in the TG and the cornea of miR-183C KO vs WT control mice^27^. This increased CX3CL1 in the cornea may contribute to the increased number of corneal resident myeloid cells in naïve miR-183C conventional KO^51^ as well as the SNS-CKO vs their corresponding WT control mice that we reported earlier^27^.

Furthermore, we identified that the genes encoding both TLR4 and FPR1 are direct targets of miR-183C. Inactivation of miR-183C results in an upregulation of TLR4 and FPR1^27^, and contributes to the enhanced response to PA bacteria. These data suggest that miR-183C regulates PA-induced production/secretion of CX3CL1 and sP by targeting the genes encoding the receptors for the bacteria on the TG sensory neurons (**Fig.6**).

Collectively, our data demonstrate that miR-183C imposes a multitude of regulation on TG sensory neurons modulating their basal production and secretion of functionally important chemokine and neuropeptide to their target tissues and their response to PA infection (**Fig.6**).

These data elucidate a potential mechanism underlying the role of corneal sensory innervation in the initiation and development of PA keratitis. However, one of the caveats is that, in the current study, the TG sensory neurons are derived from entire TG ganglion, rather than cornea-specific sensory neurons. Therefore, the observations may not fully represent the functions of cornea-specific sensory neurons. Additional studies using cornea-specific sensory neurons are required to fully validate their functions in PA keratitis. In addition, our study brings about numerous new questions. For example, what are the mechanisms of the release of these neuropeptides under homeostasis and/or upon PA bacterial infection? Are the neuropeptides released simply by enzymatic processing, or by synaptic vesicles (SV) or extracellular vesicles (EV), which have been shown to play important roles in intercellular communication and PA keratitis^110^? Are CX3CL1 and sP released in the same vesicles or are they sorted and released by different mechanisms? Are the production and secretion of other neuropeptides, e.g., CGRP, also regulated by miR-183C? In addition of neuropeptides, do the miRNAs of miR-183C from the TG sensory neurons also released to the corneal niche through SV and/or EV? Will these miRNAs enter other constituent cells in the corneal niche and modulate their functions and response to PA infection, and therefore, modulate PA keratitis? Furthermore, our data showing that PA induced TG sensory neurons production/secretion of key chemokine and neuropeptide suggest that PA bacteria activate TG nociceptor sensory neurons through TLR4 and FPR1, and, therefore, may cause acute neurogenic pain, in addition to inflammatory pain^71, 111, 112^. Since our data also demonstrated that miR-183C regulates PA-induced neuropeptide production/ secretion, we speculate that miR-183C may modulate PA-induced ocular pain, which is one of the major symptoms of PA keratitis^113^. Answers to these questions will provide greater precision of the mechanism underlying the roles of miR-183C in TG sensory neurons modulating the corneal niche and the pathogenesis of PA keratitis. Such knowledge will be the prerequisite for development of miRNA-based therapy for the treatment/prevention of PA keratitis and management of PA keratitis-associated ocular pain. Besides PA keratitis, our discoveries in this report have important implications to sensory neuron-bacteria interaction and neuroimmune interactions in the pathogenesis of other bacterial infectious diseases.

## Acknowledgement

This work is supported by grants from the National Eye Institute, National Institutes of Health (R01EY026059 to SX; R01EY035231,and P30EY004068 to LDH, U01EY034711 to AJS, R01EY026891 to AJS); a Research to Prevent Blindness unrestricted grant to the Department of Ophthalmology, Visual and Anatomical Science, Wayne State University School of Medicine. We thank Joseph Stephen Flot, research technician of the Department of Ophthalmology, University of Pittsburgh, for teaching us to make the microfluidic chambers; and Manoranjan Santra, research technician of the P30 Vision Core of the Department of Ophthalmology, Visual and Anatomical Sciences for his assistance in mouse genotyping.

---

**Fig. S1.**
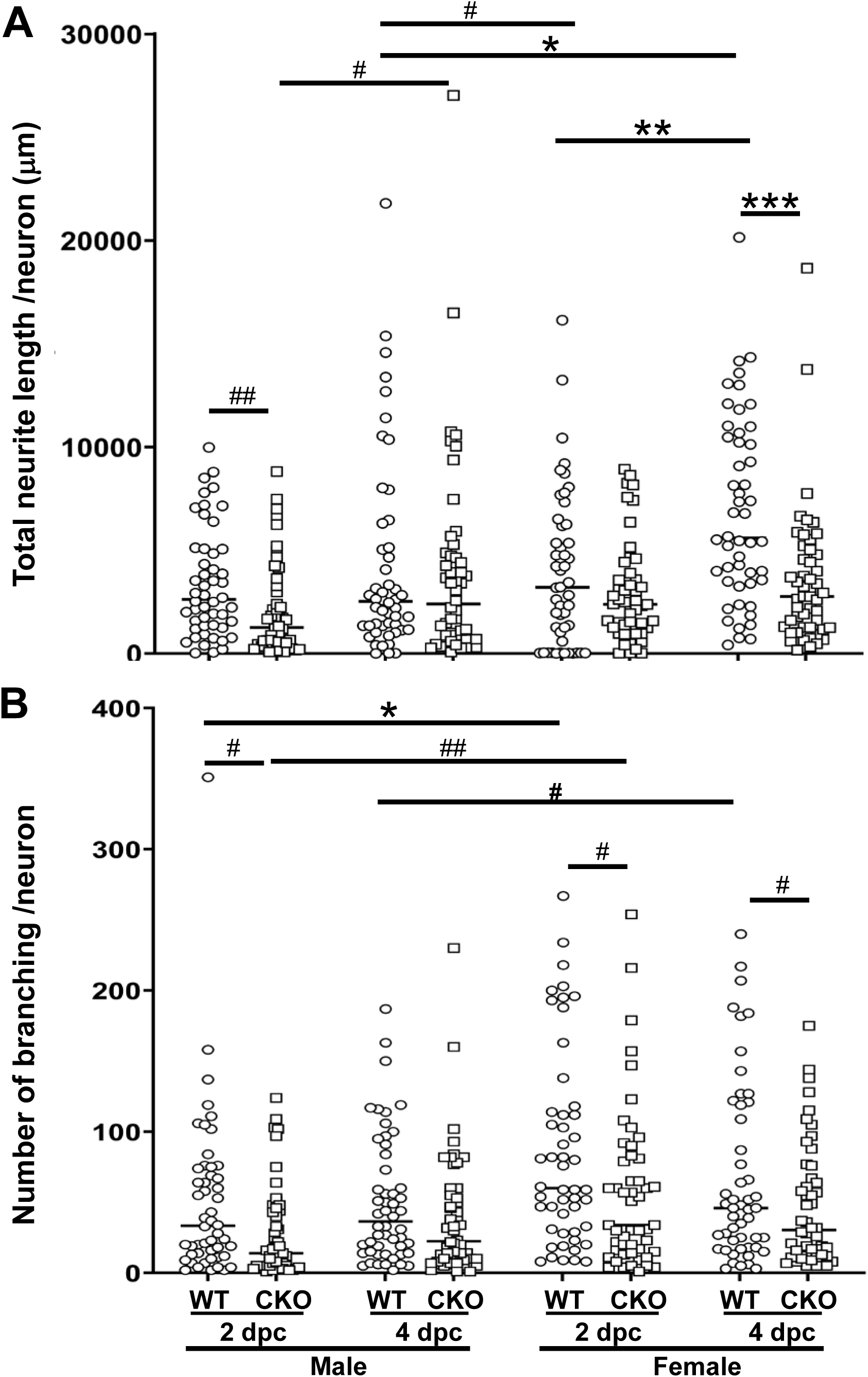
Comparison of neurite length (A) and branching (B) in male vs female mice by neurite tracing assay in 24-well plates. *: p<0.05, **: p<0.01, ***:p<0.001 by one-way ANOVA of multiple sample comparison with Bonferroni post hoc test. #: p<0.05, ##: p<0.01 by Student’s *t* test.

**Fig. S2.**
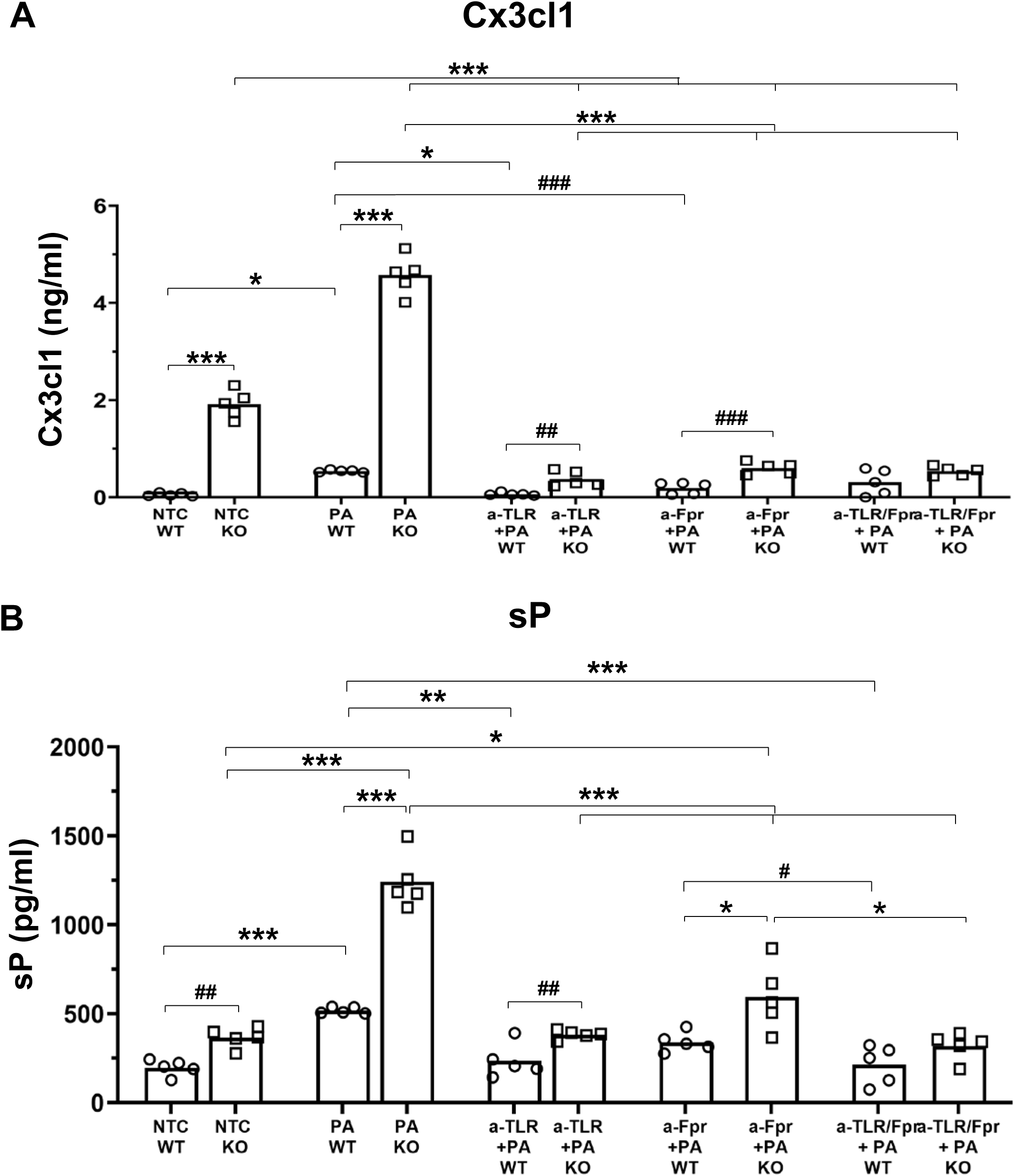
TLR4 and Fpr1 mediate PA-induced expression of neuropeptides, Cx3cl1 and substance P (sP). ELISA assays in the cell lysates of the soma chambers. NTC: no treatment control; a-TLR+PA: axon chambers pretreated with TLR4-specific antagonist, LPS-RS Ultrapure (20 µg/mL) 2 hours before PA infection; a-Fpr+PA:, the axon chambers pre-treated with Fpr1 antagonist, Boc-MLF (2 µg/mL) 2 hours before PA infection; a-TLR/Fpr+PA: axon chambers pre-treated with LPS-RS Ultrapure (20 µg/mL) and Boc-MLF (2 µg/mL) 2 hours before PA infection. *: p<0.05, **: p<0.01, ***:p<0.001 by one-way ANOVA of multiple sample comparison with Bonferroni post hoc test. #: p<0.05, ##: p<0.01, ###: p<0.001 by Student’s *t* test.

**Fig. S3.**
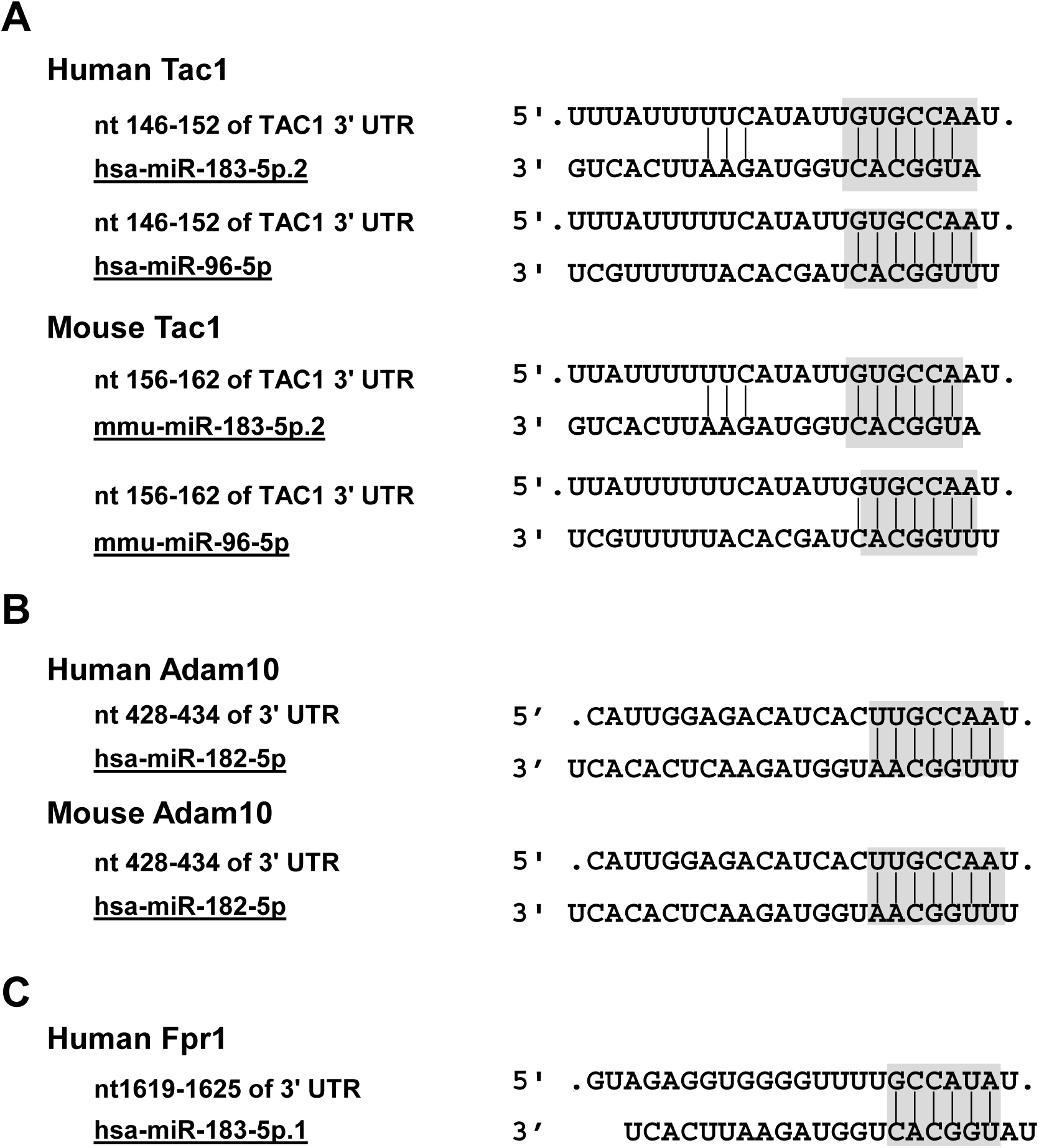
Tac1, Adam10 and Fpr1 are predicted targets of miR-183C based on TargetScan algorithm. Sequence alignment of miR-182, miR-183, and/or miR-96 with their target sites in human and/or mouse Tac1 (A), Adam10 (B) and Fpr1 (C). The shaded sequences are seed sequences of the designated miRNAs and their target sites.

## Notes

### Competing Interest Statement

The authors have declared no competing interest.

